# Single-cell analysis of the human pancreas in type 2 diabetes using multi-spectral imaging mass cytometry

**DOI:** 10.1101/2021.03.29.437504

**Authors:** Minghui Wu, Michelle Y.Y. Lee, Daniel Traum, Jonathan Schug, Irina Kusmartseva, Mark A. Atkinson, HPAP Consortium, Klaus H. Kaestner

## Abstract

Type 2 diabetes mellitus (T2D) is a chronic age-related disorder characterized by hyperglycemia due to the failure of pancreatic beta cells to compensate for increased insulin demand, typically associated with peripheral insulin resistance. However, despite decades of research, the pathogenic mechanisms underlying T2D remain poorly defined. Imaging mass cytometry (IMC) enables multiplexed assessment of the abundance and subcellular localization of multiple proteins on the same tissue section. Herein, we utilized IMC with a panel of 34 antibodies to quantify key markers of pancreatic exocrine, islet, and immune cells as well as stromal components. We employed this panel to analyze over 2.1 million cells from 16 pancreata obtained from donors with T2D and 13 pancreata from age similar non-diabetic controls. In the T2D pancreata, we observed significant alterations in islet architecture, endocrine cell composition, and surprisingly immune cell constituents. Thus, both HLA-DR positive CD8 T cells and macrophages were enriched intra-islet in the T2D pancreas. These efforts demonstrate the utility of IMC to investigate complex events at the cellular level in order to provide new insights into the pathophysiology of T2D.

## INTRODUCTION

The endocrine pancreas, although constituting only 1 to 2 percent of the organ’s mass, is highly effective in regulating blood glucose in a narrow physiological range (Atkinson et al., 2020; In’t Veld and Marichal, 2010). The insulin-producing beta cells, which comprise 20 to 60% of islet cells in the normal human pancreas (Wang et al., 2019), are able to adapt extensively to metabolic demand. However, this capacity is not without limits, with type 2 diabetes (T2D) ensuing when insulin-secretion becomes insufficient (DeFronzo et al., 2015). A seminal study by Deng and colleagues analyzed pancreatic islet structure and function in a large cohort of T2D cases (Deng et al., 2004) and noted that islet mass from the T2D pancreas was not only reduced (i.e., 50% of that of controls), but also reported that these islets were smaller and enriched for glucagon-producing alpha cells. Importantly, insulin secretion was not only impaired in terms of maximal output, but also in glucose threshold, which shifted from 7 to 12 mM glucose (Deng et al., 2004).

Two potential mechanisms are frequently invoked to account for the beta cell failure observed with T2D, namely beta cell ‘exhaustion’ due to prolonged elevated insulin production and the associated ER stress (Bilekova et al., 2020; Kataoka and Noguchi, 2013), or a loss of beta cell mass through apoptosis following long-term exposure to high glucose/high lipid levels through a process termed ‘glucolipotoxicity’. The latter mechanism is proposed to involve the Toll-Like Receptor (TLR)-mediated sensing of palmitate in beta cells, followed by chemokine production and recruitment of macrophages to the islet (Eguchi and Nagai, 2017). Indeed, islet inflammation has increasingly been proposed as the key player in beta cell dysfunction in T2D (reviewed in Eguchi and Nagai, 2017). Experimental support for this model derives from both rodent studies (Eguchi et al., 2012; Masters et al., 2010; Westwell-Roper et al., 2014) as well as analyses of human pancreatic samples (Ehses et al., 2007; Kamata et al., 2014; Richardson et al., 2009). For example, an increased frequency of islet resident macrophages was observed in leptin-deficient mice that become diabetic as a consequence of extreme hyperphagia (Ehses et al., 2007). Importantly, there was a shift from a predominantly M1 to M2 macrophage phenotype in this animal model (Cucak et al., 2014). Similarly, an elevated abundance of macrophages was also observed in human islets obtained from T2D patients, both with and without amyloid deposition (Ehses et al., 2007; Richardson et al., 2009). The relevance of this observation to T2D pathogenesis in humans has been supported by clinical studies utilizing interleukin 1 beta (IL-1β) receptor antagonists or neutralizing antibodies to IL-1β (Cavelti-Weder et al., 2012; Larsen et al., 2007). However, from these studies it cannot be excluded that the observed improvements in glycemia and insulin secretion were the result of drug effects on peripheral tissues such as skeletal muscle, liver or adipose tissue. Perhaps most importantly, these studies, while provocative and hypothesis-generating in terms of their ability to address why T2D develops, they none-the-less remain limited in their interpretation due to constraints in terms of complex data analysis and the technologies required to generate such information.

Indeed, determining the pathogenic events that occur in the T2D pancreas is critical not only for an improved understanding of the disorder’s pathogenesis but in addition, the development of novel therapeutic methods. However, the histopathological analysis of the human pancreas has been severely hampered by the fact that the pancreas cannot be easily or safely biopsied (Atkinson, 2014; Mueller et al., 1988). Thus, the vast majority of investigations have focused on rodent models such as the aforementioned leptin-deficient mouse or animals fed with various high fat/high sucrose diets in order to induce obesity and/or insulin resistance. While the rodent models have the tremendous advantage of accessibility and ease of genetic manipulation, it is likely that they do not capture all aspects of islet failure in human T2D. In response to this, high quality human pancreata, obtained from deceased organ donors, are increasingly being procured in limited number through organizations such as the Network for Pancreatic Organ donors with Diabetes (nPOD) (Campbell-Thompson et al., 2012) and more recently, the NIH supported Human Pancreas Analysis Program (HPAP; (Kaestner et al., 2019).

With these recent increases in tissue availability, opportunities exist to obtain information regarding islet cell subtyping, architectural frameworks, immune cell composition, the identification of cell-cell interactions, as well as other cellular phenotypes that might contribute to the pathogenesis of T2D. Recent advances in combinatorial immunolabeling and multiplexed protein detection *in situ* have allowed for proteomic information that is not simply additive, but rather offer crucial insights into the cellular states and complex biological functions executed by different cell types within the islet environment. Here, in order to obtain such a multiplexed view of all critical cell types and states in the T2D pancreas, we have utilized imaging mass cytometry (IMC) technology, which we have previously employed successfully to analyze the pathogenesis of type 1 diabetes (T1D) (Wang et al., 2019). IMC applies metal-conjugated antibodies to label tissue sections which are subsequently ablated by a UV-laser spot-by-spot. The resulting particle plumes are then transferred to a mass spectrometer for signal detection and quantification (Giesen et al., 2014). The precise laser registration in the sample ablation step allows for signal detection at 1 μm resolution. More importantly, the discrete time-of-flight measurement of isotope mass in the mass spectrometry instrument facilitates multiplexing. In the following, we utilized a panel of 34 antibodies to analyze multiple pancreata from both control and T2D patients and document striking changes in islet composition, immune cell infiltration, and cell to cell contacts within the islet microenvironment.

## RESULTS AND DISCUSSION

### Study overview

We employed IMC for the co-registered determination of relative protein levels for 34 antigens relevant to human pancreas biology, including markers of all endocrine cell types (C-peptide of insulin, glucagon, somatostatin, PP and ghrelin), those specific for pancreatic ductal and acinar cells (Pan-Keratin, CD44, PDX1, Carbonic anhydrase II and pS6), endothelial and stromal cells (CD31 and Nestin), extracellular matrix (Collagen Type 1), key transcription factors (PDX1, NKX6.1), immune cells of the myeloid as well as lymphoid lineages (CD45; CD20, CD3, CD4, CD8, CD45RO, CD68, CD14, CD11b, HLA-DR, CD56, CD57 and Granzyme B [GnzB]), as well as MHC class I and II proteins (HLA-ABC and HLA-DR) (See Table S1). After validation of our staining protocol using HPAP tissue samples, we selected our experimental cohorts from the nPOD archive to include 13 non-diabetic (ND) and 16 T2D cases (Table S2). Importantly, the two cohorts did not differ by sex distribution, age or BMI (Table S2). Because of regional differences in islet composition and, possibly, disease presentation, we analyzed sections from the head, body and tail of the pancreas separately, wherever available (Figure 1A). Figure 1 B, C presents three representative multi-color overlays each for a non-diabetic and a T2D pancreas obtained from the same IMC experiment. The selected images illustrate some of the major observations we quantitate below, namely the relative loss of beta and gain of alpha cells, the increase in type 1 collagen deposition, and changes in the immune cell compartment in the T2D pancreas.

**Figure 1.**
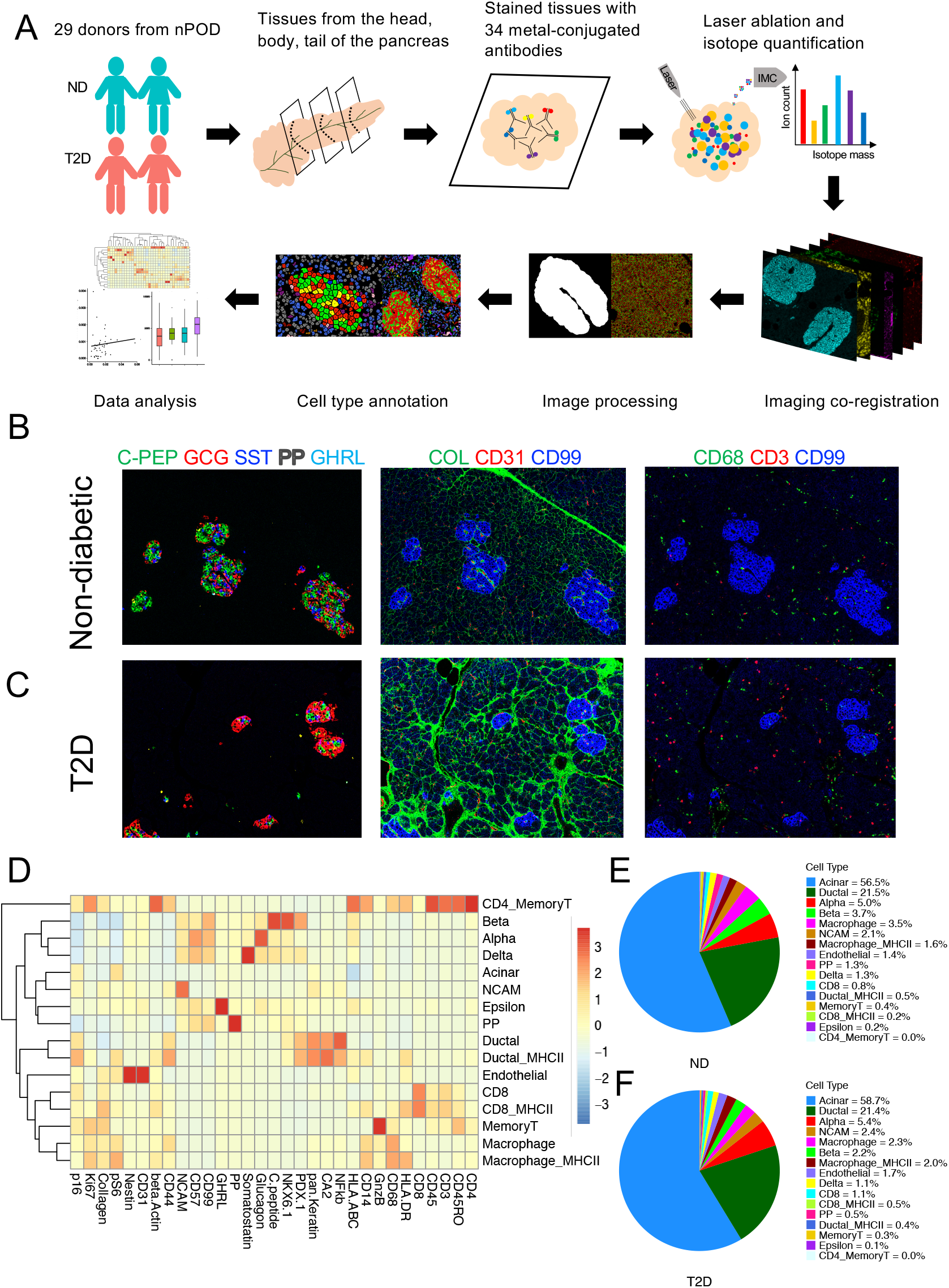
Imaging Mass Cytometry Study of the Human Pancreas Overview. See also Table S1–S2 and Figure S1–3. (A) Imaging mass cytometry workflow (B, C) Representative multiple channel image overlays of non-diabetic (B) and T2D (C) pancreas: left to right displayed channels are as follows: C-peptide (green), Glucagon (red), Somatostatin (blue), PP (white), and Ghrelin (cyan); Collagen (green), CD31 (red), CD99 (blue); CD68 (green), CD3 (red), CD99 (blue). (A) Heat map of mean expression values of each protein in each cell type. Color indicates Z score. (E, F) The proportion of each cell type in non-diabetic (E) and T2D (F).

### Image Data Processing and Cell Type Profiling

To allow for quantitative analyses, we performed image processing to produce a table of cells with expression levels for the 34 antigens plus the two DNA channels in each region of interest (ROI) imaged, while retaining the location information for each cell (see STAR methods). We then performed cell type calling to annotate the identity of each single cell. As the first step of image processing, we performed pixel-level processing to remove machine-induced artifacts such as streaks of extreme intensities and single “hot pixels”. We then segmented each raw image into islets and individual cells, based on the strong pan-endocrine CD99 staining and DNA plus membrane staining, respectively. The expression of each protein was summarized by the mean pixel intensity across the segmented cell mask, turning the segmented cell images into a cell-protein expression table. We then performed cell-level compensation to mitigate any low-level spillovers observed between a given heavy metal isotope detection channel, its neighboring channels, and the channels containing potential oxidation products. Finally, the cells were annotated through two rounds of clustering, with the first round separating cells into the three major cell types of endocrine, immune, and ‘other’, while the second round assigned specific cell subtypes (see STAR methods).

### Overall cell type composition of the human pancreas

Through our IMC analysis of pancreatic tissue sections from these 29 donors, we were able to assign cell types to 2,100,853 cells while retaining their positional information and relative protein expression levels. The heatmap in Figure 1D illustrates results from the cell type annotation efforts. For example, cells with high levels of C-peptide, NKX6.1, PDX1 and CD99 were identified as beta cells, while cells expressing both Nestin and CD31 were defined as endothelial cells. For macrophages, CD8 T cells and pancreatic ductal cells, we noted two subpopulations differing in their expression levels of HLA-DR, an MHC class II molecule (Figure 1D); we therefore considered them as two distinct cell states throughout the analyses.

Our approach of selecting ROIs based on the presence of at least one islet imparts a relative bias toward the endocrine compartment in our analysis, which reached approximately 11% of total cells analyzed (alpha, beta, delta, epsilon and PP cells combined). Taking this into account, the overall composition of the pancreas was not dramatically different between the control and T2D groups, with 78.6±8.7% of the ROIs comprised of exocrine (ductal and acinar) cells (Figure 1E,F). However, a subsequent, more detailed analysis of the alpha and beta cell density demonstrated that the frequency of beta cells was significantly decreased in the T2D pancreas (Figure 2A,C), with the strongest effect seen within the body of the organ (Figure 2A, *P* < 0.01), with the head and tail of the pancreas was affected to a lesser extent (Figure 2A). The tissue density of alpha cells was reciprocally increased in the T2D pancreas, again with the strongest impact in the body (Figure 2B). Next, we analyzed the alpha to beta cell ratio, an effort which demonstrated that it was significantly increased in the diabetic state (Figure 2C). Endocrine cell proportions by donor and region are quantified in Figure S1.

**Figure 2.**
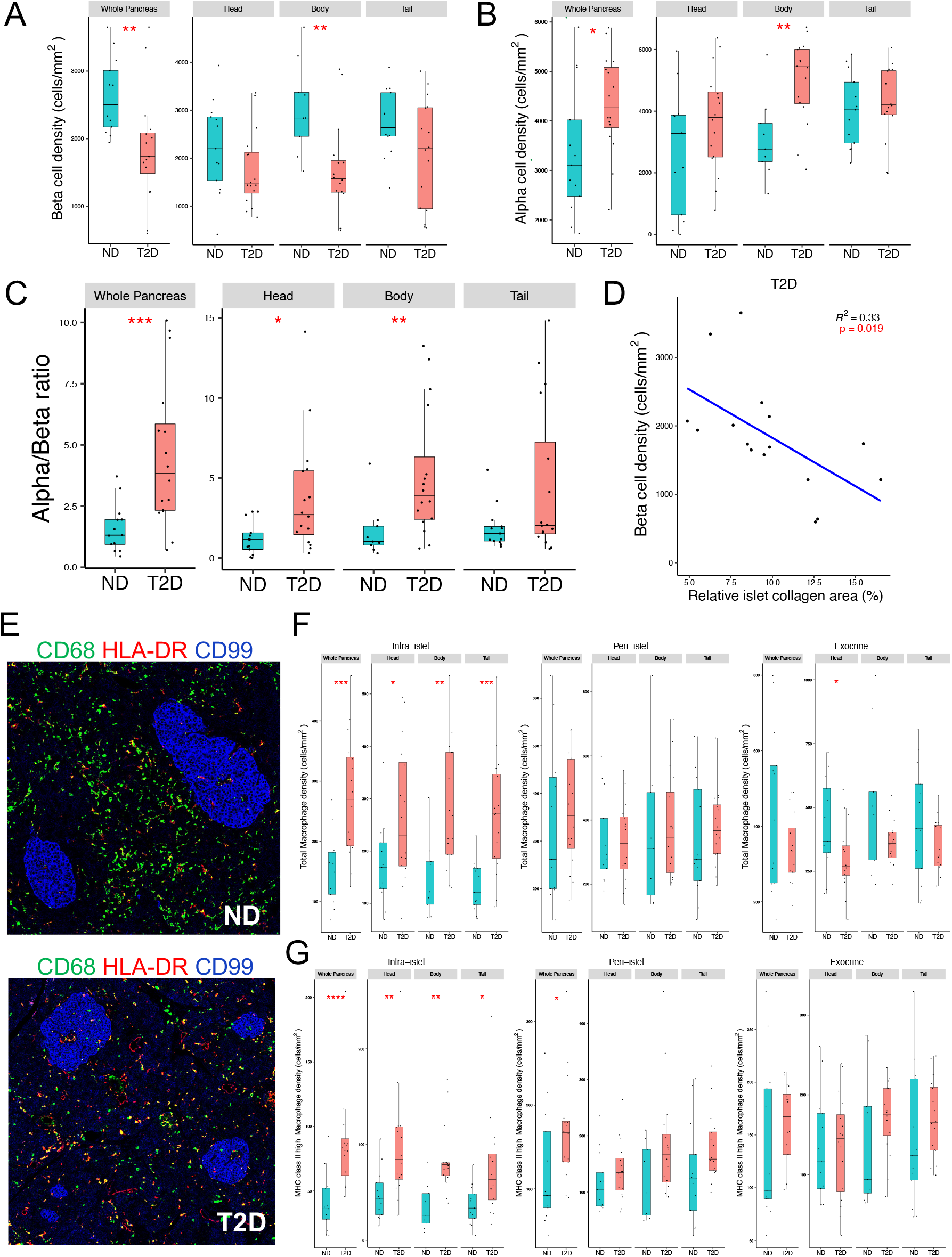
Altered alpha and beta cell as well as macrophage distribution in the T2D pancreas. (A) Boxplots showing the density of beta cells in the human pancreas, analogous to panel A. *p < 0.05, **p < 0.01. Mann-Whitney U test. (B) Boxplots showing the density of alpha cells in the human pancreas, either for the whole organ or separately for the head, body or tail. Box plots display median, 25-75% range and outliers. *p < 0.05, **p < 0.01. Mann-Whitney U test. (C) Boxplots of the alpha cell/beta cell ratio in the ND and T2D pancreas. **p < 0.05, **p < 0.01, ***p < 0.001. Mann-Whitney U test. (D) Beta cell density is inversely proportional to type 1 collagen deposition in the T2D pancreas. Islet collagen is measured by the percentage of expanded islet area (islet region + within 50μm from islet boundary) with positive collagen signal. Statistical significance tested by linear regression t-test. Each dot demonstrates one donor. (E) Representative image of CD68^+^ macrophages with or without HLA-DR expression in the pancreas from non-diabetic (above) and T2D (below) organ donor. Islets outlined by CD99 (blue). (F) Boxplots showing the total macrophage density intra-islet, peri-islet or in the exocrine pancreas. Data are presented for the whole pancreas and separately for head, body and tail of the organ. ND, non-diabetic; T2D, type 2 diabetic. *p < 0.05, **p < 0.01. Mann-Whitney U test. (G) Boxplots showing HLA-DR^high^ macrophage density intra-islet, peri-islet or in the exocrine pancreas. Data are presented for the whole pancreas and separately for head, body and tail of the organ. ND, non-diabetic; T2D, type 2 diabetic. **p < 0.05, **p < 0.01, ***p < 0.001, ****p < 0.0001. Mann-Whitney U test.

Changes in the T2D pancreas have been reported to be related to the duration of the disease (Chen et al., 2017); therefore, we analyzed our data to assess this possibility. Indeed, we observed a significant positive correlation between alpha cell density and T2D duration in the entire organ as well as in the body of the pancreas, along with a significant reduction in beta cell density that was notably associated with disease duration as reflected in the pancreas body (Supplemental Figure 2).

While the replication rates of human endocrine cells in adulthood are exceedingly low, they are higher in alpha cells than in beta or delta cells throughout life (Wang et al., 2019). To assess whether altered proliferation rates contribute to the shift in alpha and beta cell densities in T2D, we assessed their positivity for Ki67. In the ND pancreas, we observed the expected linear decrease in replication rates between 14 and 65 years of age, with an approximate three-fold higher rate for alpha cells (Figure S3). In contrast, in the T2D tissues, proliferation rates of both alpha and beta cells were low even in younger individuals (Figure S3).

As noted earlier, we observed an increase in type I collagen deposition in the extracellular matrix of the T2D pancreas (Figure 1B,C). Therefore, we asked whether type 1 collagen deposition, a classical sign of fibrosis, is related to beta cell abundance. While there was no correlation between the two parameters in the ND pancreas (data not shown), collagen density was significantly anti-correlated with beta cell density in T2D individuals (Figure 2D), suggesting a link between the fibrotic process and remaining beta cell mass.

### Intra-islet HLA class II expressing macrophages are increased in T2D

Next, we turned our attention to the immune cell compartment within the pancreas. CD68^+^ macrophages are present at highest density within the exocrine pancreas, demonstrated an abundance of approximately 400 cells per mm^2^, which did not differ overall between the T2D and ND samples, except in the head region (Figure 2F). However, when probing the positional information retained by IMC to quantify intra-islet macrophages, we observed a highly significant, nearly two-fold increase in their abundance in tissues from T2D individuals, which was present in all three regions of the organ analyzed (Figure 2F). As mentioned previously, macrophages in the human pancreas can be divided into HLA class II high and low cells based on levels of HLA-DR. While the HLA-DR^high^ cells constituted between 15%-65% of the total macrophage population in the pancreas (depending on donor and area), their relative abundance was increased throughout the organ in the T2D pancreas, with the greatest increase of approximately 2.1 fold intra-islet (Figure 2G).

### CD8 T cells are increased in the T2D pancreas

While the primary cause of T1D is autoimmunity, this is not the case for T2D. Nevertheless, islet inflammation has long been proposed to contribute to beta cell impairment in T2D (Ehses et al., 2007), but with sparse description within the literature. Therefore, we analyzed the abundance and spatial distribution of CD8 T cells in the head, body, and tail of the pancreas in detail (Figure 3). We observed that the density of CD8 T cells was increased in the exocrine pancreas of T2D donors, with trends in all three regions of the organ, but were only significant in the body (Figure 3B). Notably, the fold-change difference in CD8 T cell abundance between control and T2D was highest when we considered the intra- and peri-islet regions of the tissue. We also detected a small subset of CD8 T cells that stained positive for HLA-DR. Expression of MHC class II molecules on CD8 T cells is commonly employed as a marker of activation (Ndhlovu et al., 2015). Interestingly, we observed that HLA-DR^high^ CD8 T cells were increased significantly in the T2D pancreas, elevated up to 2.9 T2D fold in the intraislet region (Figure 3C).

**Figure 3.**
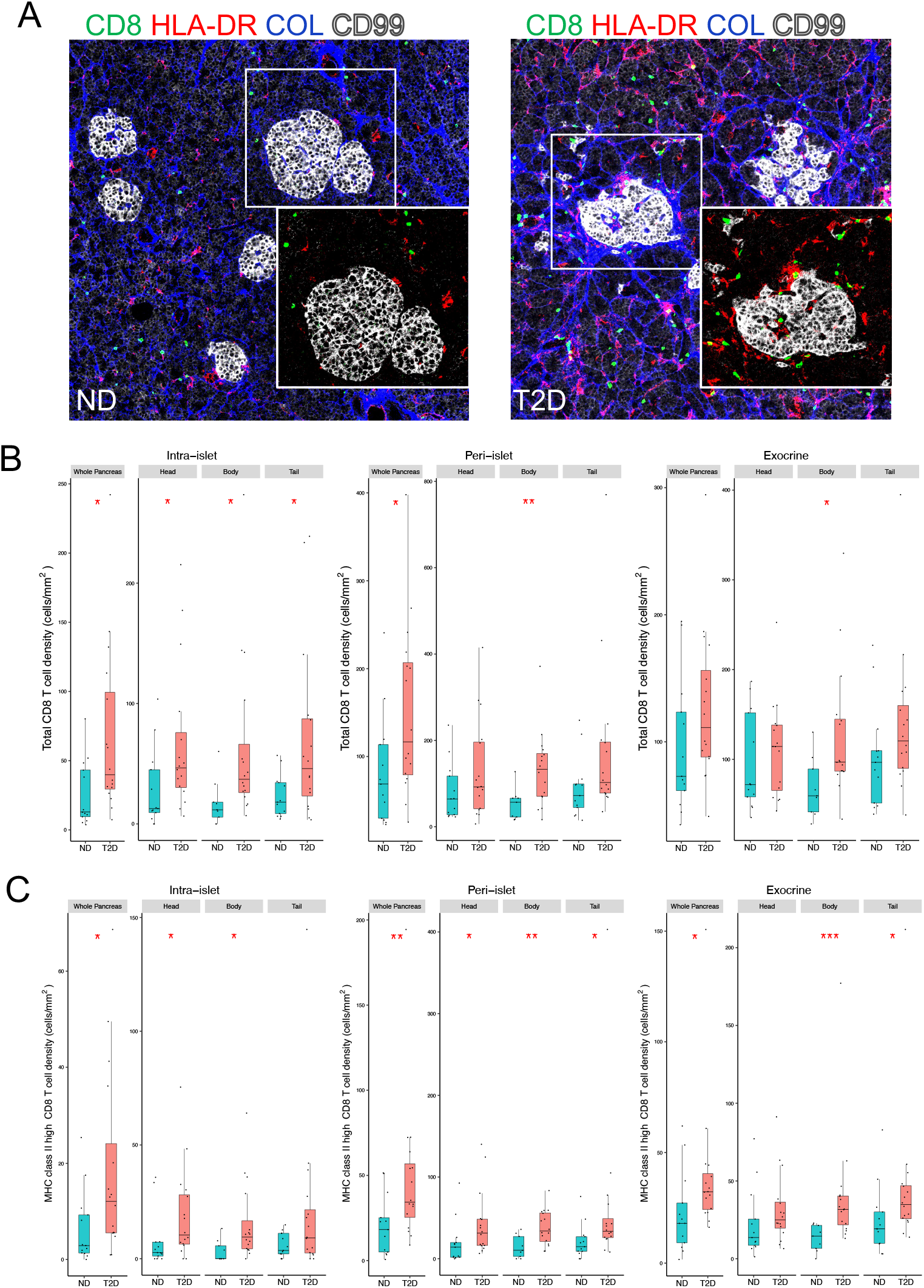
Intra-islet CD8 T and CD8/HLA-DR^high^ T cell density is increased in T2D. CD8 T cell and HLA class II expression in the pancreas of non-diabetic (left) and T2D (right) organ donors. CD68 (green), HLA-DR (red), Collagen (blue), and CD99 (white). (A) Boxplots showing total CD8 T cell density intra-islet, peri-islet or in the exocrine pancreas. Data are presented for the whole pancreas and separately for head, body and tail of the organ. ND, non-diabetic; T2D, type 2 diabetic. *p < 0.05. Mann-Whitney U test. (B) Boxplots showing CD8/ HLA-DR^high^ T cell density intra-islet, peri-islet or in the exocrine pancreas. Data are presented for the whole pancreas and separately for head, body and tail of the organ. ND, non-diabetic; T2D, type 2 diabetic. *p < 0.05, **p < 0.01. Mann-Whitney U test.

### Neighborhood analysis of IMC data

The retention of spatial information following segmentation and cell type identification in IMC enabled neighborhood analyses for the more than 2 million cells captured in our study. Figure 4A illustrates the accuracy of our cell type annotation pipeline. On the left of this figure, we show an overlay of an ROI with seven markers that are expressed in the major epithelial cells of the pancreas, while the image on the right displays the 14 cell types identified after our cell type profiling. Having established the high degree of similarity between the actual image and post-segmentation rendering, we proceeded to perform HistoCAT neighborhood analysis to examine the frequencies of specific cell-cell interactions (Schapiro et al., 2017) as schematized in Figure 4B and FigS5. Each cell type was considered as the ‘query cell,’ and the frequency of every other cell type being in its neighborhood was recorded. Next, we compared the frequency with a null distribution and calculated an enrichment score representing the deviation from the null (see STAR methods). Following this, we compared the neighborhood enrichment between ND and T2D donors, as summarized in the heatmap in Figure 4C. Shown on the left are the labels of the query cells (row labels) and on the bottom, the labels of the neighborhood cell types. Plotted are the differences in specific cell-cell interactions. The shading of each square indicates whether this interaction is enriched in the pancreas of ND or T2D donors, and black boxes highlight those demonstrating statistical significance. Through this analysis we determined that CD8 T cells and CD68^+^ macrophages are more likely to be in contact with islet beta cells, while macrophages are less likely to be near acinar cells in T2D.

**Figure 4.**
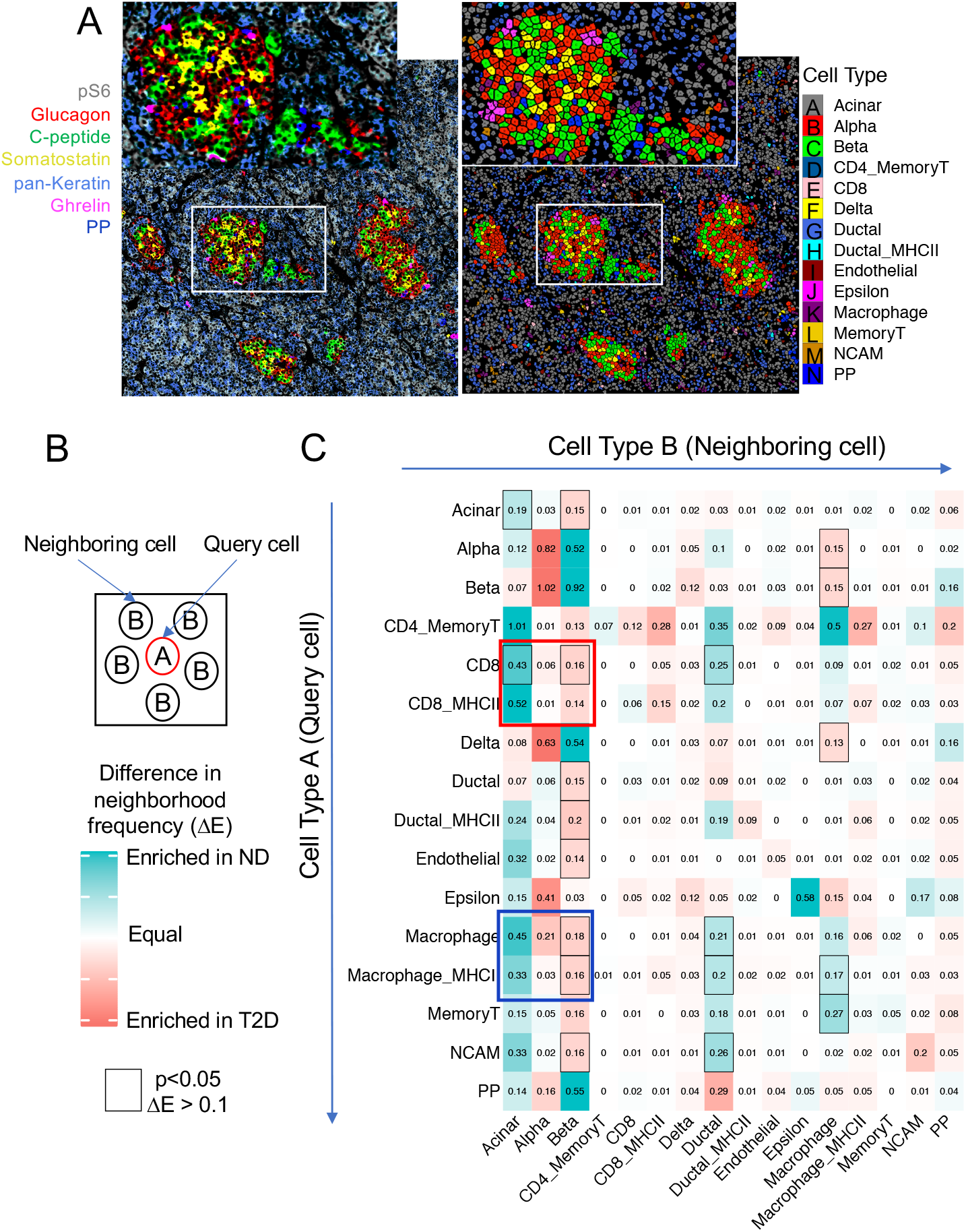
Neighborhood analysis. See also Figure S4. (A) Cell type annotation of the 14 major cell types determined after image segmentation was projected back to assemble the tissue map shown on the right. Note the close match to the original IMC image shown on the left, confirming the cell type annotation pipeline. (B) Principle of neighborhood analysis. For any cell “A”, termed query cell, the cell types of the adjacent cells (neighboring cells, “B”) were counted and summarized as a frequency matrix. Enrichment scores were calculated by comparing the observed frequency to those obtained from 100 permutations. Enrichment scores were then compared for statistical significance between control and T2D by permutation-test. (C) Heat map displaying cell–cell interaction frequencies between non-diabetic and T2D donors. Statically significant results are outlined by a black box. Teal color indicates interactions that are more frequent in non-diabetic (ND) and red those that are more abundant in the T2D donors. The red box highlights the finding that the presence of CD8 T cells near beta cells is more likely in the T2D pancreas, while the blue box emphasizes the point that macrophage/beta cell interactions are more likely in T2D compared to the ND pancreas.

In sum, this effort established the power of IMC for the analysis of the histopathological changes in human T2D, and with this, brought novel insights regarding the pancreas in T2D; efforts, that provide important clues into the disorder’s pathogenesis. We have confirmed the relative loss of beta cells and increase in alpha cells in T2D, a facet of the disease phenotype which is dependent on disease duration. We discovered that the ageing-dependent decline in islet endocrine cell proliferation is accelerated in individuals with T2D (Figure S3). These findings suggest that the relative increase in alpha cell number in T2D is not driven by higher proliferation rates, which leaves trans-differentiation from another cell type or *de novo* generation from adult endocrine progenitor cells yet to be defined, as the remaining likely options. Finally, our efforts also noted a negative correlation between islet collagen area and beta cell density, providing further evidence that fibrosis might play a contributing role in beta cell loss in T2D.

An increased abundance of islet-resident macrophages was first reported by the Donath group in 2007, with *in vitro* experiments suggesting that islets incubated with very high glucose and palmitate concentrations could secrete pro-inflammatory cytokines and chemokines, which could in turn stimulate monocyte/macrophage recruitment (Ehses et al., 2007). Herein, we confirmed and extend these findings by demonstrating not only that intra-islet macrophage density is increased in T2D, but that a subset of these expressed high levels of HLA-DR, the significance of which awaits further analysis. The presence of CD8 T cells in the T2D pancreas has also been reported previously, but with the conclusion that CD8 T cells are only increased in the exocrine pancreas but not in islets (Rodriguez-Calvo et al., 2014). In addition, when employing FACS analysis on isolated islets, Butcher and colleagues found no change in CD3 T cells abundance in T2D (Butcher et al., 2014), although it is possible that this approach missed leukocytes that migrated out of the islet during isolation. In contrast, we observed increased density of CD8 T cells within islets, and this relative shift of CD8 T cells from the exocrine pancreas to the islet in T2D was confirmed by our neighborhood analysis (Figure 4B). Interestingly a subset of islet-resident CD8 T cells were positive for HLA-DR, a common marker of T cell activation (Ndhlovu et al., 2015), and the density of these islet resident CD8^+^HLA-DR^+^ cells in was also increased in T2D. These findings suggest that activated CD8 T cells could contribute to islet inflammation in human T2D.

#### Limitations of Study

We would note that there are several potential limitations to our study. While we were able to analyze a sufficient number of cases to define statistically significant differences between non-diabetic and T2D donors in terms of islet composition, immune cell infiltration and cellular neighborhoods, our efforts were unfortunately underpowered to analyze heterogeneity among the T2D donors. Disease heterogeneity among the clinical presentation of T2D patients is increasing recognized, with several classification systems recently proposed (Ahlqvist et al., 2020). Future studies should include organ donors typed for future consensus subclassifications of T2D to study this issue further.

Beyond this, there were additional limitations related to the IMC platform. While this technology enables the simultaneous detection of several dozen antigens and is not plagued by issues of tissue auto-fluorescence often seen in immunofluorescence approaches, IMC can result in low sensitivity for some proteins since “exposure time” cannot be increased, as can be performed for fluorescence-based imaging platforms. In addition, subcellular location analysis is limited because each pixel has 1 μm squared dimensions. Finally, IMC destroys the tissue in the laser ablation process, and therefore additional orthogonal experiments cannot be performed.

## SUPPMEMENTAL INFORMATION

Supplemental Information includes three tables and four figures.

## ACKNOWLEDGEMENTS

We thank Dr. Mingyao Li (University of Pennsylvania) for participating and contributing to the discussion about IMC analysis. Funding was provided by NIH grants U01 DK123594 (to KHK) and P01 AI42288 (to TMB and MAA). This research was performed with the support of the Network for Pancreatic Organ donors with Diabetes (nPOD; RRID:SCR_014641), a collaborative type 1 diabetes research project sponsored by JDRF (nPOD: 5-SRA-2018-557-Q-R) and The Leona M. & Harry B. Helmsley Charitable Trust (Grant#2018PG-T1D053).

## AUTHOR CONTRIBUTIONS

MW and DT performed imaging mass cytometry experiments, MW, MYYL, and JS performed computational and statistical data analysis, IK, MAA and KHK wrote and edited the manuscript, KHK supervised the study.

## CONTACT FOR REAGENT AND RESOURCE SHARING

Further information and requests for resources and reagents should be directed to and will be fulfilled by the Lead Contact, Klaus Kaestner.

## STAR METHODS

### EXPERIMENTAL MODEL AND SUBJECT DETAILS

The pancreas tissue samples were obtained from the Network for Pancreatic Organ Donors with Diabetes (nPOD, https://www.jdrfnpod.org) under IRB approval by the University of Florida.

Donor clinical information was obtained from nPOD and T2D diagnosis followed ADA guidelines.

### METHOD DETAILS

#### Antibody Panel and labeling

20 of 34 conjugated antibodies were purchased from FluidigmR (https://www.fluidigm.com). The other 14 unconjugated antibodies were obtained from different vendors (Table S1) and conjugated to lanthanide metals using the MaxPar X8 Multimetal Labeling Kit (Fluidigm) according to the manufacturer’s protocol. Antibodies were diluted with 0.5% BSA in PBS.

#### Tissue staining and image acquisition

Five to eight μm FFPE pancreas sections were stained with a cocktail of 34 antibodies (Table S1). Tissues were de-paraffined with xylene for 30 minutes and rehydrated in the sequential ethanol from 100% to 70% with changes every 5 minutes. After transfer to ddH_2_O for 5 minutes, we performed epitope retrieval in a decloaking chamber with HIER buffer (10mM Tris, 1mM EDTA, pH9.2) for 30 minutes at 95°C. Tissue sections were allowed to cool to room temperature in HIER buffer and then transferred to PBS for 20 minutes. After blocking in 3% BSA for 1 hour, the tissues were stained with the antibody cocktail at 4°C overnight. The next day, the tissues were labeled by 1:400 dilution of Ir-intercalator solution (Fluidigm 201192B) in PBS for 30 minutes to label nuclei. Slides were washed in PBS two times for 5 minutes, dipped 2 minutes in ddH_2_O and airdried before IMC acquisition.

Following Fluidigm’s operation instruction and daily tuning we acquired the IMC images at a laser frequency of 200 Hz using Fluidigm’s Hyperion instrument. 1,000 μm x 1,000 μm regions around islets were selected based on analysis of adjacent H/E stained sections. Finally, we converted the mcd files to tiff images using Fluidigm’s MCD viewer.

#### Image processing

We captured a total of 260 images (regions of interest of 1,000 μm x 1,000 μm) from our cohort of 16 T2D and 13 non-diabetic donors. Prior to signal quantification, streaks of pixels with extreme intensity were detected based on two background channels (Iridium 113 and Iridium 115) and were removed using an approach adopted from Wang et al. (Wang et al., 2019). Specifically, a 5×5 grid was used to search for pixels with intensity among the 2% of the whole image and greater than 2x median intensity of the grid (the median was calculated excluding the center row). A list of pixels fulfilling this requirement in the first two background channels were tested again in the following protein channels and pixels with value greater than 5x the median intensity of the 5×5 grid, were replaced with the median intensity. Lastly, single “hot pixels” with intensity greater than 50 (intensity units) of the maximum value in its 3×3 neighborhood were replaced with the local maximum, using a CellProfiler plugin (Zanotelli et al., 2020).

#### Cell Segmentation

Cells masks were generated following a published pipeline (Schapiro et al., 2017). Firstly, CD99 was used to detect cell membranes of endocrine and exocrine cells. Due to the large difference in CD99 intensity between the two cell populations, the ‘Enhance Local Contrast (CLAHE)’ function in Fiji was applied to CD99 channel (blocksize=39, histogram=256, maximum=40, mask=’none’). Secondly, four channels were selected for cell segmentation: CD45, CD68, enhanced CD99, Iridium 193 (the Iridium DNA intercalator labels nuclei). Images were enlarged by 2-fold and converted to the h5 format. H5 files were then imported into Ilastik v1.3.3 (Berg et al., 2019) for pixel classification training. Specifically, a Random Forest model was trained using the 37 default features to classify pixels into one of three classes: nuclei, membrane, or background. Four images were labelled manually for training, then the classifier was applied onto the rest of images. The outputs of Ilastik were three probability maps, one for each class. The nuclei probability map was used to detect primary object (nuclei) in CellProfiler with the minimum cross entropy method. Primary objects with an area of less than 5 pixels were filtered out. Secondary objects were identified using the Distance – B method. Specifically, primary objects recognized in the previous step were expanded with guidance of the cell membrane probability map, with maximum expansion constrained as 10 pixels. Resulting cell masks were resized by 0.5-fold to the original scale and cells with area less than 25 pixels were removed. Lastly, the cell masks were used to measure mean protein expression for each channel and each cell in the original IMC images post pre-processing steps. The x and y coordinate of all cells in the image were also recorded.

#### Spillover compensation

Signal crosstalk between channels were compensated using the functions from CATALYST R package (v.1.12.2) (Chevrier et al., 2018) based on the isotope purity matrix provided by Fluidigm.

#### Islet Segmentation and measurement

We employed a similar workflow for islet segmentation. Specifically, a separate model was trained to generate islet probability maps based on the CD99 channel. The probability maps were segmented into the islet masks using CellProfiler using a similar process as described for cell masks. We determined islet related measurements with CellProfiler as well. Specifically, for islet collagen percentage, we expanded the islet mask by 50μm, segmented collagen positive regions, and quantified the percentage of expanded islet area positive for collagen signal.

### QUANTIFICATION AND STATISTICAL ANALYSIS

#### Data transformation and normalization

We normalized mean pixel intensity of each target channel for each cell as described below before downstream quantitative analyses. Firstly, we transformed the raw mean pixel intensity data by log_2_ (mean intensity+1). For most antibody targets, the most common cell signal value corresponds to cells that do not express the target. Thus, for each image, we identify the most frequent signal value mode, S_0_, using the highest peak in a smoothed estimate of the distribution of the log transformed signal values (S) for each channel (R function density [S, n=2^16]). Subsequently, we calculated the normalized value S_n_ = S – S_0_ and set S_n_ < 0 to 0. Lastly, S_n_ is pooled across all images and clipped to the 99.99th percentile value for each channel to remove any outliers.

#### Clustering and meta clustering

We performed clustering following the FlowSOM workflow (Van Gassen et al., 2015). Cells with normalized protein mean expression were clustered in two steps. First, we grouped all cells into 225 clusters using a self-organizing map. Markers used for clustering in this stage were selected based on a high signal-to-noise ratio and their information content for cell type calling as follows: C-peptide, Glucagon, Somatostatin, PP, PDX.1, NKX6.1, Ghrelin, CD99, Carbonic Anhydrase 2, NF-κb, CD44, Nestin, CD31, CD56, CD57, HLA.DR, β-Actin, HLA.ABC, phospho-S6(pS6), Foxp3, p16, CD8, CD3, CD45RO, CD4, CD45, Granzyme B, CD68, CD14, CD20. Next, we combined the 225 groups into 40 clusters using the ‘MetaClustering_consensus’ function in the FlowSOM package. Next, the 40 clusters were grouped manually into 3 major cell types: ‘endocrine’, ‘immune’, and ‘other’ based on the mean and distribution of the marker proteins. In the second step, each cell type was reclustered using the same workflow as in the first step. Cell were divided into 225 groups with a self-organizing map and recombined into 50 clusters. Markers used were the same with the first stage except that we excluded CD99 for endocrine clusters. The resulting 150 clusters, 50 for each major cell type were annotated based on the distribution and mean of protein expression. For ambiguous clusters, images were inspected to determine the most appropriate cell type on the basis of cell location and morphology.

#### Neighborhood analysis

We adopted the HistoCAT neighborhood analysis to examine cell-cell interaction difference between conditions (Schapiro et al., 2017). We used the R implementation available on Github, “BodenmillerGroup/neighbouRhood”. Cells were considered as neighbors if their centers were less than 20 pixels apart. For each ROI, we calculated an interaction score between two cell types, with one cell type (A) being the query cell and the other cell type (B) being the neighboring cell. Specifically, I_AB_ is the average number of cell type B (neighboring cell) in the neighborhood of cell type A (center cell). Then, a permutation was employed by shuffling cell labels and thus resulted in a null distribution, assuming that all cells interacted randomly. We used 100 rounds of permutation and calculated a mean for each type of interaction. We then calculated an enrichment score, the difference between observed interaction and expected interaction under the null model. Lastly, we used permutation tests with 5,000 iterations to compare the enrichment score between T2D and ND, and Benjamini-Hochberg adjustment for multiple testing was applied to the resulting p-values. An interaction was accepted as significantly enriched in one condition if the adjusted p-value < 0.05. In all the images, the boundary cells were excluded from the neighborhood calculation because they do not represent a true neighborhood due to image cutoff.

#### Statistical methods

For Figure 2A-C,2F, 2G, 3B and 3C, Mann-Whitney test with p ≤ 0.05 was used for statistical testing. For data associated with Figures 2D, S2A, S2B, S3A, S3B, statistical analyses were performed with fit linear model. For Figure 4B, Benjamini-Hochberg adjustment for multiple testing was applied. P-values were represented as follows: *p < 0.05; **, p <0.01; ***, p<0.001, **** p<0.0001.

### DATA AND SOFTWARE AVAILABILITY

Raw image data will be available to download from the nPOD website: https://www.jdrfnpod.org.

---

**Supplemental Figure 1.**
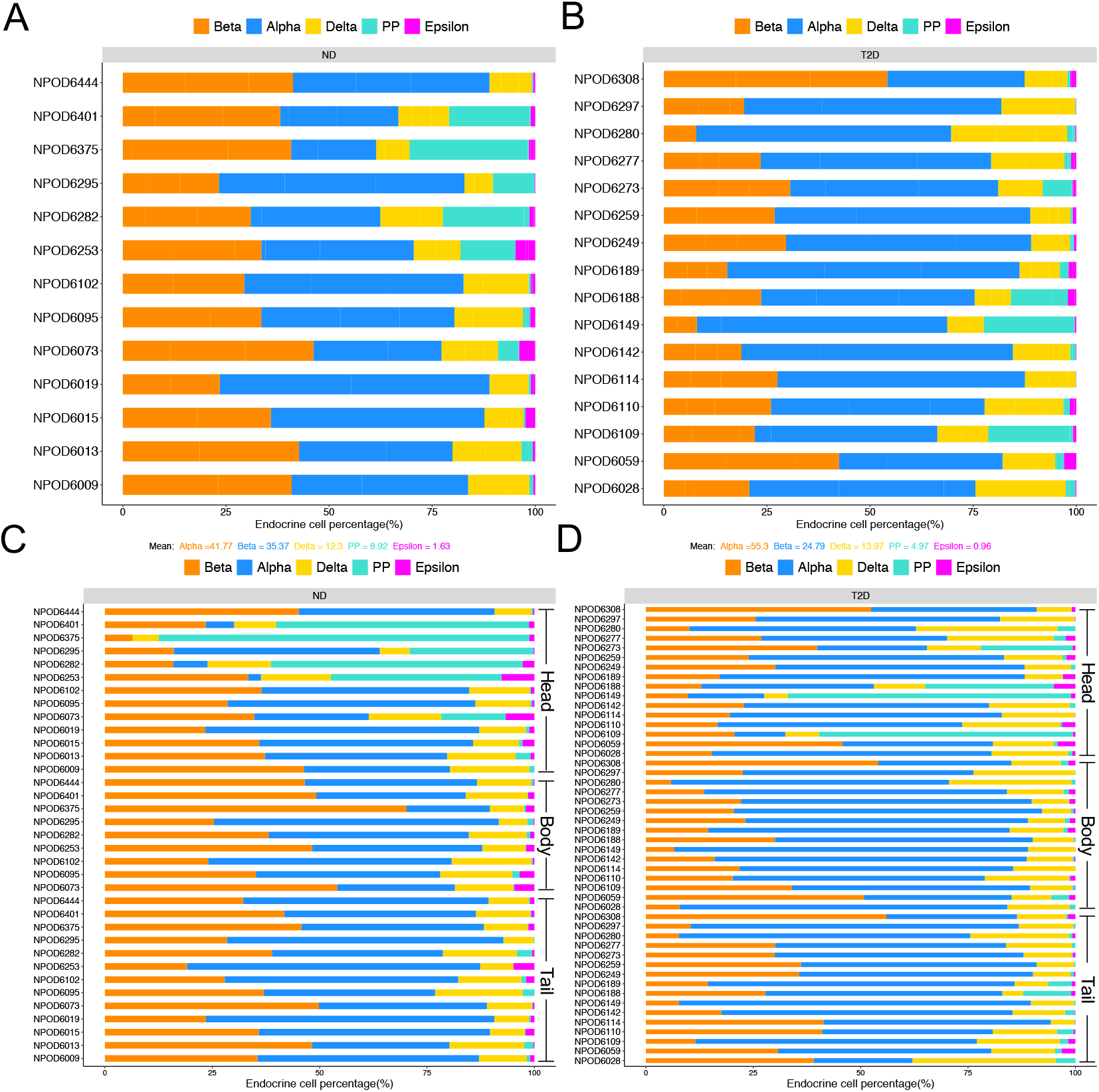
Endocrine cell composition in the human pancreas as determined by IMC. (A, B) Stacked bar plot showing endocrine cell percentage of each donor. (A) non-diabetic and (B) T2D. (C, D) Stacked bar plot showing endocrine cell percentage in each region of each donor of no-diabetic (C) and T2D (D).

**Supplemental Figure 2:**
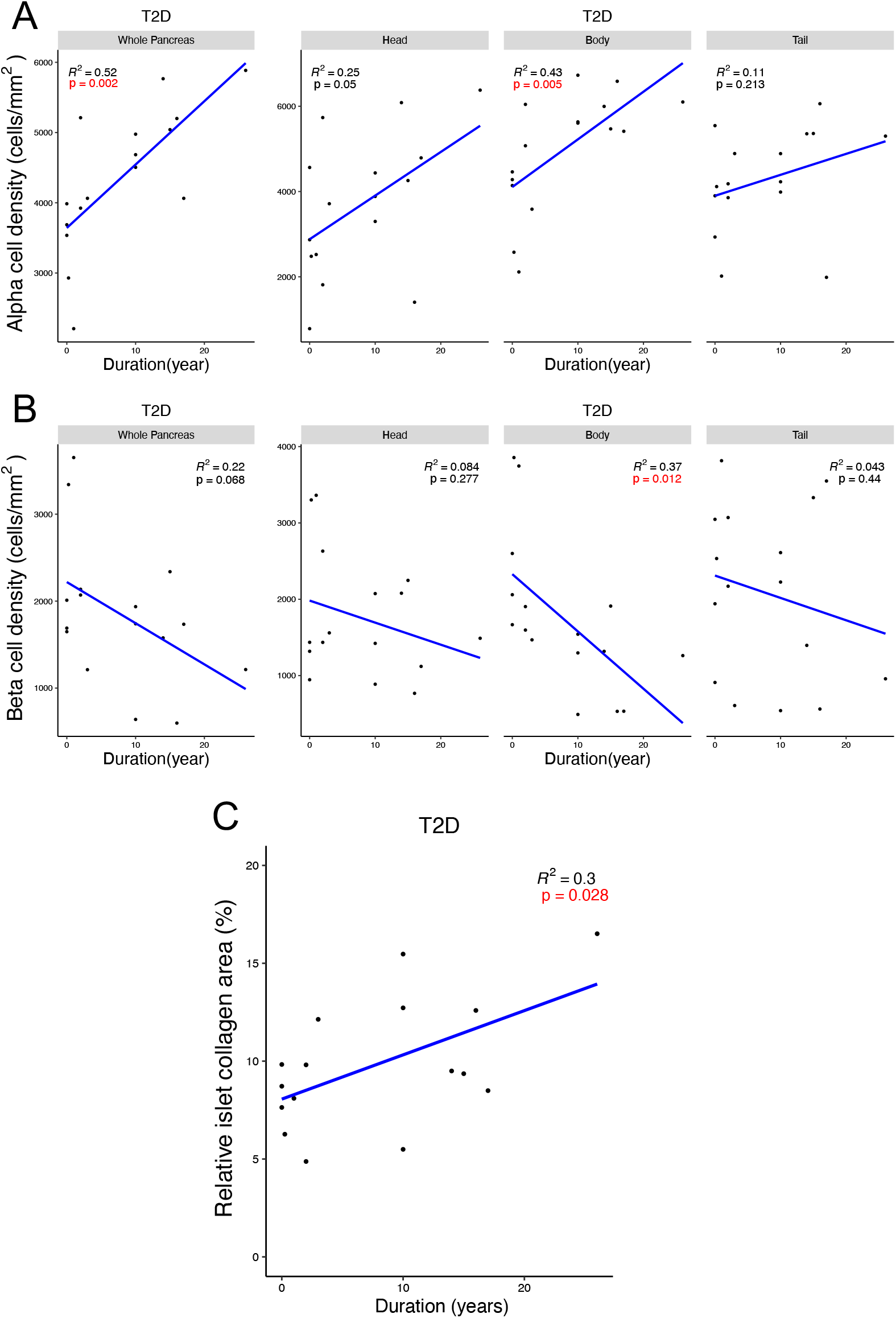
Alpha and beta cell density in the T2D pancreas is related to duration of disease. (A, B) Scatter plot showing the correlation between alpha cell (A), beta cell (B) density and the duration of disease in T2D organ donors. Linear regression was used to determine statistical significance.

**Supplemental Figure 3.**
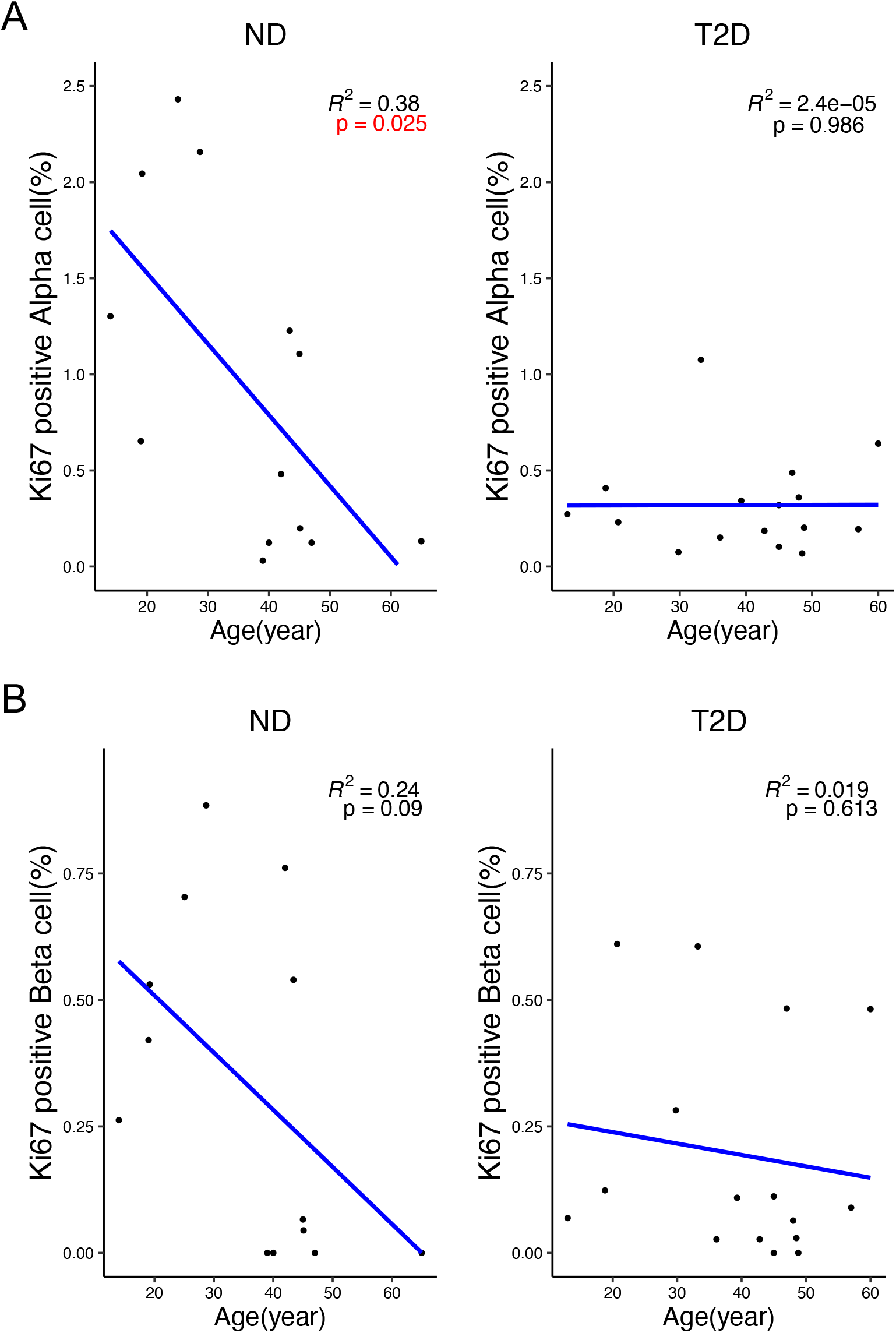
Quantification of proliferating alpha and beta cells with respect to age in non-diabetic and T2D organ donors. (A, B) Correlation between the percentage of Ki67-positive alpha cell (A), beta cell(B) and age of non-diabetic (left) and T2D (right) organ donors. Linear regression was used to determine statistical significance.

**Supplemental Figure 4.**
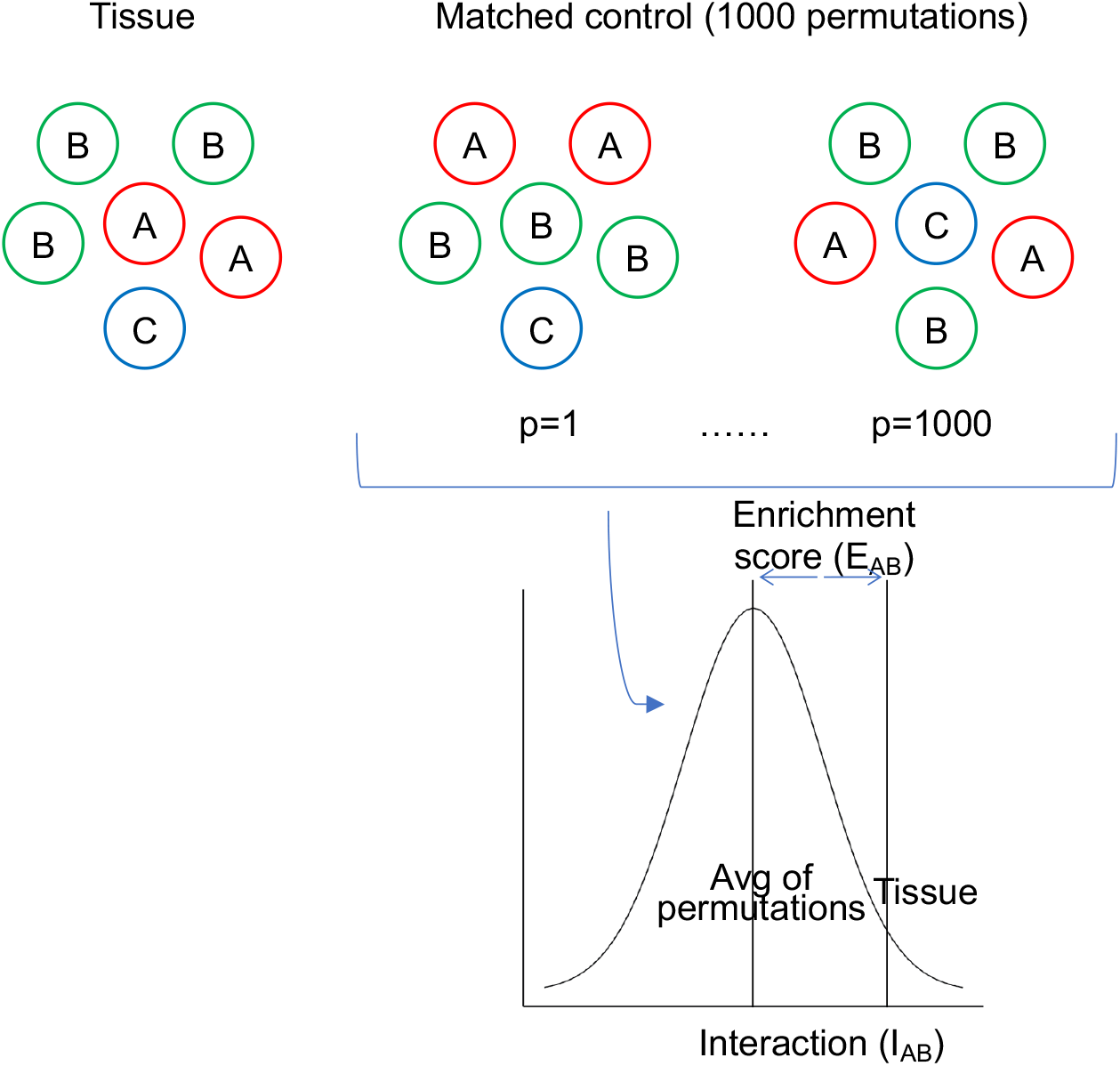
Illustration of the enrichment score calculation used for the neighborhood analysis. (A) The frequency of cell-cell interaction is compared with a null distribution by permuting the cell type labels. For each cell-cell interaction, A being the query cell and B being the neighboring cell in question, I_AB_, is calculated. Next, the labels of all the neighboring cells are shuffled for 500 times and an average interaction of the permutations is calculated. Lastly, the average I_AB_ of permutated distribution is subtracted from the observed I_AB_ in tissue to obtain an enrichment score (E_AB_) for interaction between A and B.

## KEY RESOURCES TABLE

| REAGENT or RESOURCE | SOURCE | IDENTIFIER |
| --- | --- | --- |
| Antibodies |  |  |
| 141Pr-HLA-ABC (Clone EMR8-5) | BD Biosciences | Cat#565292 |
| 142Nd-CD57 (Clone HCD57) | Fluidigm | Cat#3142007B |
| 143Nd-CD31 (Clone C31.3+C31.7+C31.10) | LifeSpan BioSciences | Cat#LS-C390863-100 |
| 144Nd-CD14 (Clone EPR3653) | Fluidigm | Cat#3144025D |
| 145Nd-C-peptide (Clone 3A1) | Thermo | Cat#MA1-22710; RRID: AB_558517 |
| 146Nd-Nestin (Clone 196908) | Fluidigm | Cat#3146015B |
| 147Sm-Glucagon (Clone C-11) | Santa Cruz | Cat#sc-514592; RRID: AB_2629431 |
| 148Nd-Pan-Keratin (Clone C11) | Fluidigm | Cat#3148020D |
| 149Sm-CD11b (Clone EPR1344) | Fluidigm | Cat#3149028D |
| 150Nd-CD44 (Clone IM7) | Fluidigm | Cat#3150018B |
| 151Eu-PDX1 | R&D systems | Cat#AF2419; RRID: AB_355257 |
| 152Sm-CD45 (Clone 2B11) | Fluidigm | Cat#3152016D |
| 153Eu-CD56 | Proteintech | Cat#14255-1-AP; RRID: AB_2149421 |
| 154Sm-Beta-actin (Clone 2F1-1) | Fluidigm | Cat#3154021D |
| 155Gd-FoxP3 (Clone 236A/E7) | Fluidigm | Cat#3155016D |
| 156Gd-CD4 (Clone EPR6855) | Fluidigm | Cat#3156033D |
| 158Gd-NKX6.1 | Sigma | Cat#HPA036774; RRID: AB_10673664 |
| 159Tb-CD68 (Clone KP1) | Fluidigm | Cat#3159035D |
| 160Gd-Somatostatin (G-10) | Santa Cruz | Cat#sc-13099; RRID: AB_2195930 |
| 161Dy-CD20 (Clone H1) | Fluidigm | Cat#3161029D |
| 162Dy-CD8 (Clone C8/144B) | Fluidigm | Cat#3162034D |

**Table S1.**
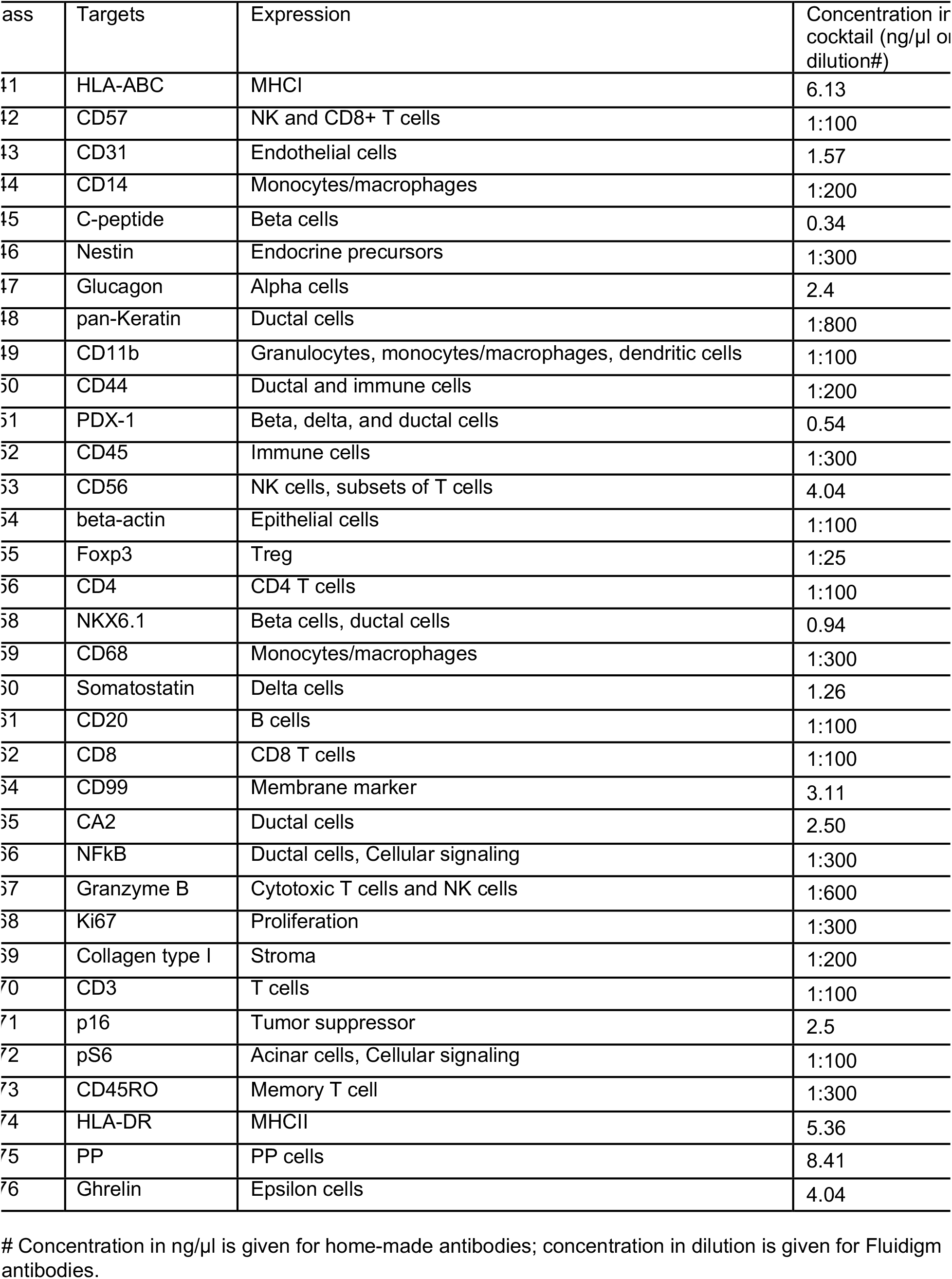
Antibodies and working concentration.

**Table S2.** Donor clinical information.

| NPOD Number | RRID | Sex | Age (years) | BMI (kg/m<sup>2</sup>) | Status |
| --- | --- | --- | --- | --- | --- |
| 6009 | nPOD, Cat# nPOD_6009, RRID:SAMN15879066 | Male | 45 | 30.6 | Non-diabetic |
| 6013 | nPOD, Cat# nPOD_6013, RRID:SAMN15879070 | Male | 65 | 24.2 | Non-diabetic |
| 6015 | nPOD, Cat# nPOD_6015, RRID:SAMN15879072 | Female | 39 | 32.2 | Non-diabetic |
| 6019 | nPOD, Cat# nPOD_6019, RRID:SAMN15879076 | Male | 42 | 31.0 | Non-diabetic |
| 6073 | nPOD, Cat# nPOD_6073, RRID:SAMN15879130 | Male | 19 | 36.0 | Non-diabetic |
| 6095 | nPOD, Cat# nPOD_6095, RRID:SAMN15879152 | Male | 40 | 35.5 | Non-diabetic |
| 6102 | nPOD, Cat# nPOD_6102, RRID:SAMN15879159 | Female | 45 | 35.1 | Non-diabetic |
| 6253 | nPOD, Cat# nPOD_6253, RRID:SAMN15879309 | Female | 19 | 34.3 | Non-diabetic |
| 6282 | nPOD, Cat# nPOD_6282, RRID:SAMN15879336 | Male | 14 | 41.9 | Non-diabetic |
| 6295 | nPOD, Cat# nPOD_6295, RRID:SAMN15879349 | Female | 47 | 30.4 | Non-diabetic |
| 6375 | nPOD, Cat# nPOD_6375, RRID:SAMN15879428 | Male | 29 | 31.8 | Non-diabetic |
| 6401 | nPOD, Cat# nPOD_6401, RRID:SAMN15879454 | Female | 25 | 31.3 | Non-diabetic |
| 6444 | nPOD, Cat# nPOD_6444, RRID:SAMN15879497 | Male | 43 | 28.4 | Non-diabetic |
| 6028 | nPOD, Cat# nPOD_6028, RRID:SAMN15879085 | Male | 33 | 30.2 | T2D_17y |
| 6059 | nPOD, Cat# nPOD_6059, RRID:SAMN15879116 | Female | 19 | 39.3 | T2D_3m |
| 6109 | nPOD, Cat# nPOD_6109, RRID:SAMN15879166 | Female | 49 | 32.5 | T2D_0 |
| 6110 | nPOD, Cat# nPOD_6110, RRID:SAMN15879167 | Female | 21 | 40.0 | T2D_0 |
| 6114 | nPOD, Cat# nPOD_6114, RRID:SAMN15879171 | Male | 43 | 31.0 | T2D_2y |
| 6142 | nPOD, Cat# nPOD_6142, RRID:SAMN15879199 | Female | 30 | 34.4 | T2D_14y |
| 6149 | nPOD, Cat# nPOD_6149, RRID:SAMN15879205 | Female | 39 | 29.1 | T2D_16y |
| 6188 | nPOD, Cat# nPOD_6188, RRID:SAMN15879244 | Male | 36 | 30.6 | T2D_0 |
| 6189 | nPOD, Cat# nPOD_6189, RRID:SAMN15879245 | Female | 49 | 36.1 | T2D_26y |
| 6249 | nPOD, Cat# nPOD_6249, RRID:SAMN15879305 | Female | 45 | 32.3 | T2D_15y |
| 6259 | nPOD, Cat# nPOD_6259, RRID:SAMN15879314 | Male | 57 | 32.3 | T2D_10y |

